# Activity of *Geranium macrorrhizum* extracts, essential oils and their main components against three relevant flagellated protozoa

**DOI:** 10.1101/2025.10.03.680318

**Authors:** Sara Marcos-Herraiz, María José Irisarri-Gutiérrez, Javier Carrión, Iris Azami Conesa, Rodrigo Suárez Lombao, Juliana Navarro-Rocha, Jose Francisco Quilez, Alejandro Fernández Barrero, Eneko Ochoa Larrigan, Azucena González-Coloma, María Teresa Gómez Muñoz, María Bailén

## Abstract

Plant-derived natural products are structurally diverse secondary metabolites that play essential ecological roles and have long been recognized for their therapeutic potential. *Geranium macrorrhizum* is known to produce bioactive extracts and essential oils rich in monoterpenoids and sesquiterpenes. While various *Geranium* species have demonstrated antiprotozoal activity, the potential of *G. macrorrhizum* against protozoan parasites remains unexplored. Given the increasing resistance of protozoan pathogens to conventional treatments, there is a pressing need to identify alternative therapeutic agents from natural sources.

Plant material was grown in greenhouse and aeroponic systems. Extracts were obtained via hydrodistillation and Soxhlet, and analyzed by GC-MS. In vitro assays were performed to evaluate antiparasitic activity and cytotoxicity. Morphological changes induced by germacrone were analyzed using light microscopy.

The study evaluated the antiparasitic potential of essential oils, extracts, and compounds from *Geranium macrorrhizum*, cultivated under controlled conditions, against *Giardia duodenalis, Trichomonas gallinae*, and *Leishmania infantum*.

Results showed broad-spectrum activity, particularly from non-polar extracts and EOs, suggesting that lipophilic compounds, especially germacrone, are key contributors to the observed effects. Germacrone exhibited strong activity and selective toxicity. Structural analogues of germacrone showed reduced efficacy, highlighting the importance of its specific chemical structure. Morphological analyses revealed parasite-specific responses to germacrone, contrasting with the more destructive effects of metronidazole and amphotericin B. The findings support *G. macrorrhizum* as a promising source of natural antiprotozoal agents and justify further investigation into its bioactive compounds and mechanisms of action.

## 1. Introduction

Plant natural products are secondary metabolites found in various parts of plants, including roots, stems, leaves, fruits and seeds. These compounds exhibit a wide structural diversity and play critical roles in mediating interactions between plants and their environment (Gervazoni et al., 2020). Natural products have been used in traditional medicine and remain a promising source for the discovery of new therapeutic agents and drugs. Over the years, numerous plant extracts, essential oils (EOs), and pure natural compounds have been identified, highlighting their immense potential in pharmaceutical and biomedical applications (Atanasov et al., 2021).

*Geranium macrorrhizum*, commonly known as Bulgarian geranium or bigroot geranium, is an herbaceous perennial species belonging to the family Geraniaceae. This species is primarily distributed across the temperate regions of the Northern Hemisphere (Aedo et al., 1998). Various species within the genus *Geranium* have historically been utilized as ornamental, medicinal, and melliferous plants. Among these, *G. macrorrhizum* holds economic significance as one of the most widely cultivated ornamental species due to its fragrance and reputed medicinal properties (Tzanova et al., 2024).

The extracts and EOs of *Geranium macrorrhizum* have demonstrated a diverse range of biological activities, highlighting its potential in various therapeutic applications. Extracts of *G. macrorrhizum* have been found to possess antimicrobial, hypotensive, spasmolytic, astringent, cardiotonic, antioxidant and sedative properties (Miliauskas et al., 2004; Radulović et al., 2012; Tzanova et al., 2024). Moreover, EOs obtained from the aerial parts of the plant, rich in monoterpenoids such as geraniol and beta-citronellol, as well as sesquiterpenes like germacrone, have shown significant insecticidal, acaricidal, antioxidant, antibacterial, and antifungal effects (Navarro-Rocha et al., 2018; Tzanova et al., 2024). These bioactive properties support the potential of *G. macrorrhizum* as a source of natural compounds for aromatherapy, phytotherapy, and potential pharmacological applications. Extracts from other *Geranium* species, such as *Geranium mexicanum* and *Geranium wallichianum*, have demonstrated antiprotozoal effects against *Giardia, Leishmania*, and *Trichomonas* (Abbasi et al., 2019; Calzada et al., 2005, 2007). However, data on the potential antiprotozoal activity of *G. macrorrhizum* extracts remain unexplored.

Protozoan parasites encompass a diverse range of eukaryotic organisms, some of which are significant contributors to globally neglected human diseases (Menezes & Tasca, 2023). These diseases pose a substantial public health challenge, affecting millions of individuals worldwide. Control and prevention strategies are increasingly undermined by the development of resistance to commonly used chemotherapeutic agents (Pramanik et al., 2019). Among the protozoan parasites of medical and veterinary importance, *Giardia duodenalis, Leishmania* spp., and *Trichomonas* spp. stand out due to their widespread impact on humans, mammals and birds.

*Leishmania* spp. cause leishmaniasis, a group of diseases transmitted through the bite of infected female phlebotomine sandflies. More than one billion people currently live in areas endemic for leishmaniasis and are at risk of infection (World Health Organization (WHO), 2024). Treatment primarily relies on pharmacological therapies; however, many available drugs (pentavalent antimonials, liposomal amphotericin B, and pentamidine) are associated with high toxicity, limited efficacy, complex administration protocols, and significant costs. Furthermore, resistant strains have emerged, complicating disease management (Sasidharan & Saudagar, 2021). *G. duodenalis* is an extracellular enteric protozoan responsible for widespread diarrheal diseases, with over 300 million cases reported annually, particularly in low-income and developing regions (Cernikova et al., 2018). Despite the availability of treatments, the increasing resistance of *Giardia* to nitroimidazole-based therapies, such as metronidazole (MTZ), poses a growing concern (Argüello-García et al., 2020; Menezes & Tasca, 2023). In birds, *Trichomonas gallinae* is a flagellated oropharyngeal parasite that causes granulomas and starvation, significantly impacting avian populations. While Columbiformes serve as the primary reservoirs, other domestic and wild bird species are also susceptible (Amin et al., 2014). The current treatment for *T. gallinae* infections also relies on nitroimidazoles, yet no preventive treatments have been approved in the EU, and therapy failures linked to resistant strains have been documented (European Commission, 1995).

There is an urgent need for alternative treatments against *Leishmania* spp., *G. duodenalis*, and *Trichomonas* spp. In this study, the antiparasitic activity of six extracts and two EOs from *G. macrorrhizum* were evaluated against *T. gallinae* and *G. duodenalis* trophozoites, as well as *L. infantum* promastigotes. Additionally, the major compounds of the tert-butyl methyl ether extract (S-TBM) were tested to assess their antiparasitic activity and cytotoxic effects. Germacrone, the primary constituent of the EOs, was further evaluated on *L. infantum* amastigotes.

## 2. Material and methods

### 2.1. Plant material

The EOs and extracts studied in this work were obtained from cultivated *G. macrorrhizum*, a plant species from the Geraniaceae family. A previous report described the field cultivation of *G. macrorrhizum* in Ejea de los Caballeros, Spain (42°8’8.73”N, 1°12’31.50”W; 346 m a.s.l.) (Navarro-Rocha et al., 2018). However, under these conditions, EOs yields were consistently low and showed considerable variability in chemical composition. Consequently, selected field-grown plants with superior biomass production were propagated for alternative cultivation means.

#### 2.1.1. Greenhouse cultivation

Rhizomes of *G. macrorrhizum* obtained from the field cultivated plants were cultivated in 1-liter containers filled with Compo Universal substrate, to which 10% (v/v) perlite was added to improve aeration and drainage. Plants were maintained for 6 months in a VENLO-type greenhouse under standard environmental conditions. Irrigation was performed by demand-driven overhead sprinkling. Fertilization was applied both in solid and liquid forms. Solid fertilization was carried out using “Compo Novatec” fertilizer with micronutrients, at a dose of 3 g per pot every six months. Liquid fertilization was conducted monthly using “Compo Huerto y Frutales” (rich in potassium), following the manufacturer’s recommended dosage. The plants flowered normally throughout this period.

#### 2.1.2. Aeroponic cultivation

The aeroponic cultivation was adapted from previously described conditions (Fraga et al., 2020). Briefly, 25 six months old greenhouse plants of 10–15 cm height were submitted to aeroponic cultivation. The plants were transferred to an aeroponic chamber (Apollo 3 system: 33 plants, 240 L, 1750 mm x 1350 mm x 750 mm) located in an environmentally controlled growth chamber (25°C) supplemented with artificial light (16:8, L:D). The roots were sprayed under constant pulverization (every 12 s) with water at 26°C supplemented with 0.2 g/L Nutrichem (20:20:20 of N:P:K—Miller Chemical and Fertilizer Crop, Hanover, PA, USA) and 0.03 % H_2_O_2_ (33% w/v Panreac Química SLU, Castellar del Vallès, Barcelona, Spain) and were cultivated for six months in water. Plant material (aerial parts and roots) was collected periodically when the length was between 10 and 15 cm. The plants did not flower under this cultivation conditions.

### 2.2 Extracts

The following extracts and EOs were obtained from the plant material by hydrodistillation (Clevenger) or Sohxlet in order to obtain compounds with different polarities (Table 1).

**Table 1.** Cultivation conditions, extraction methods, and yields of extracts and EOs.

| Production Method | Extraction Method | % Yield |
| --- | --- | --- |
| Greenhouse | Clevenger aerial (EOAP) | 0.56 |
|  | Clevenger (EOF) | 0.32 |
|  | Sohxlet Hexane (S-Hex) | 3.20 |
|  | Sohxlet TMB (S-TBM) | 1.83 |
|  | Sohxlet EtOAc (S-EtOAc) | 2.4 |
|  | Sohxlet EtOH (S-EtOH) | 2.92 |
|  | Sohxlet EtOH (D-EtOH) | 21.93 |
| Aeroponic | Sohxlet EtOH (D-EtOHa) | 29.59 |

The extraction process of EOs and extracts from *Geranium macrorrhizum* involves cultivation under two systems: greenhouse and aeroponic culture. EOs were obtained by hydrodistillation from flowers (EOF) and aerial parts (EOAP) and extracts were obtained through Sohxlet direct ethanol extraction from greenhouse-grown plants (D-EtOH) and aeroponically cultivated plants (D-EtOH), as well as through sequential Soxhlet extraction using solvents of increasing polarity: n-hexane (S-Hex), tert-butyl methyl ether (S-TBM), and ethyl acetate (S-EtOAc) (Figure 1).

**Figure 1.**
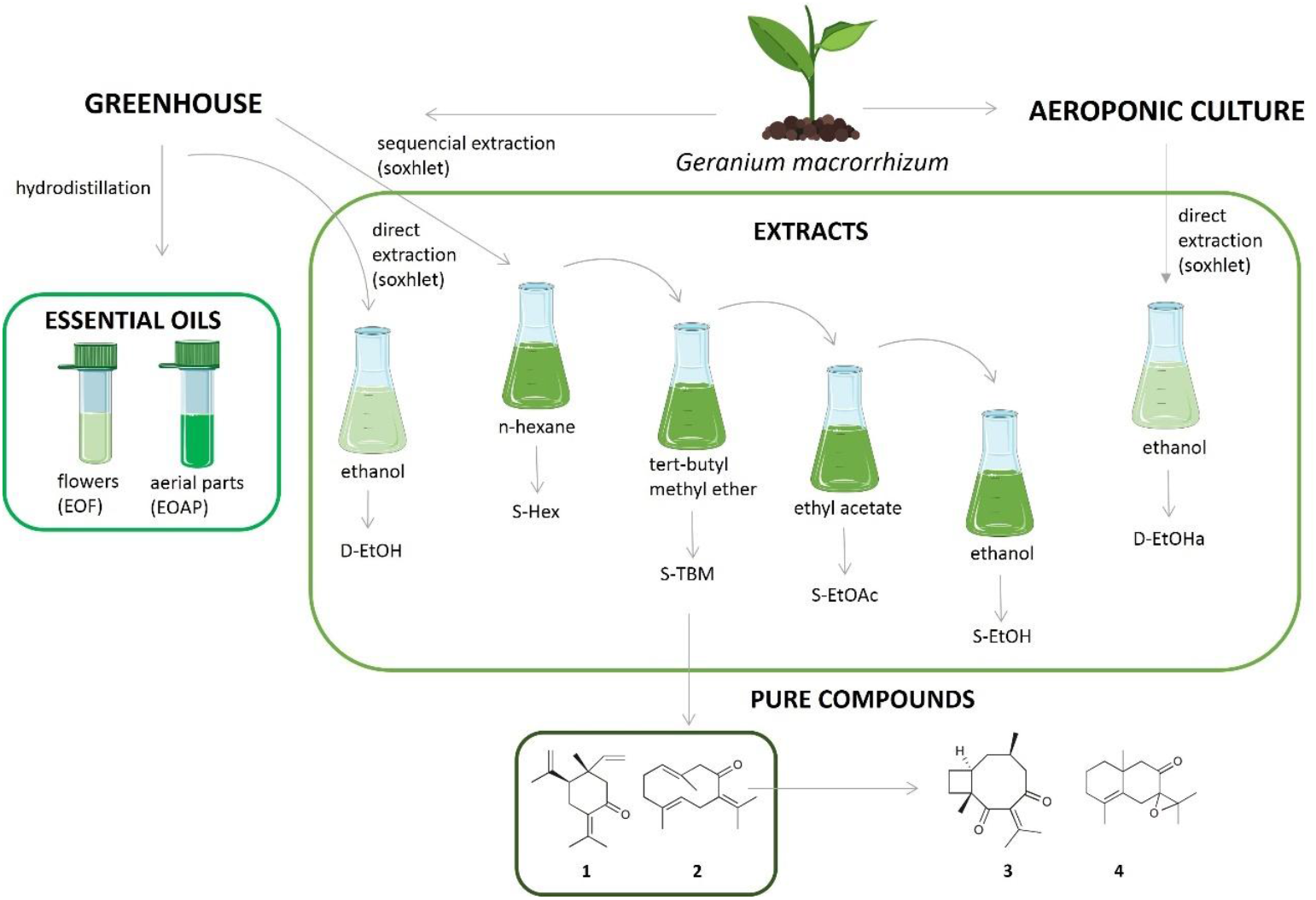
Extraction Procedures and Compound Isolation from *Geranium macrorrhizum*. Main components of S-TBM extract from *G. macrorrhizum* (greenhouse): elemenone (**1**), germacrone (**2**) and germacrone-derived compounds (**3** and **4**).

### 2.3. Extract and EOs analysis

The EOs, S-Hex, S-TBM and S-EtOAc extracts were analyzed by gas chromatography-mass spectrometry (GC-MS) using a Shimadzu GC-2010 Plus coupled to a Shimadzu GCMS-QP2010-Ultra mass detector with an electron impact ionization source at 70 eV and a Single Quadrupole analyzer and employing Helium as carrier gas. The samples were injected by an automatic injector (AOC-20i). Chromatography was carried out with a Teknokroma TRB-5 (95%) Dimethyl-(5%) diphenylpolysiloxane capillary column, 30 m x 0.25 mm ID and 0.25 μm phase thickness. The working conditions used were: Split mode injection using 1 µl of sample with a split ratio (20:1) employing a Shimadzu AOC-20i automatic injector, injector temperature 300° C, transfer line temperature connected to the mass spectrometer 250° C, and ionization source temperature 220° C. The initial column temperature was 70° C, heating up to 290° C at 6° C /min and leaving at 290° C for 15 min. All the samples (4 µg/µl) were previously dissolved in 100% dichloromethane (DCM) for injection.

The mass spectra and retention time were used to identify the compounds by comparison with those found in the Wiley database (Wiley 275 Mass Spectra Database, 2001) and NIST 17 (NIST/EPA/NIH Mass Spectral Library), while the relative area percentages of all peaks obtained in the chromatograms were used for quantification.

### 2.4. Compounds

Compounds selected for further study included the major constituents isolated from the S-TMB extract: elemenone (**1**), germacrone (**2**), and compounds **3** and **4** generated from germacrone (**2**) following protocols already reported (Galisteo Pretel et al., 2019) (Scheme 1).

**Scheme 1.**
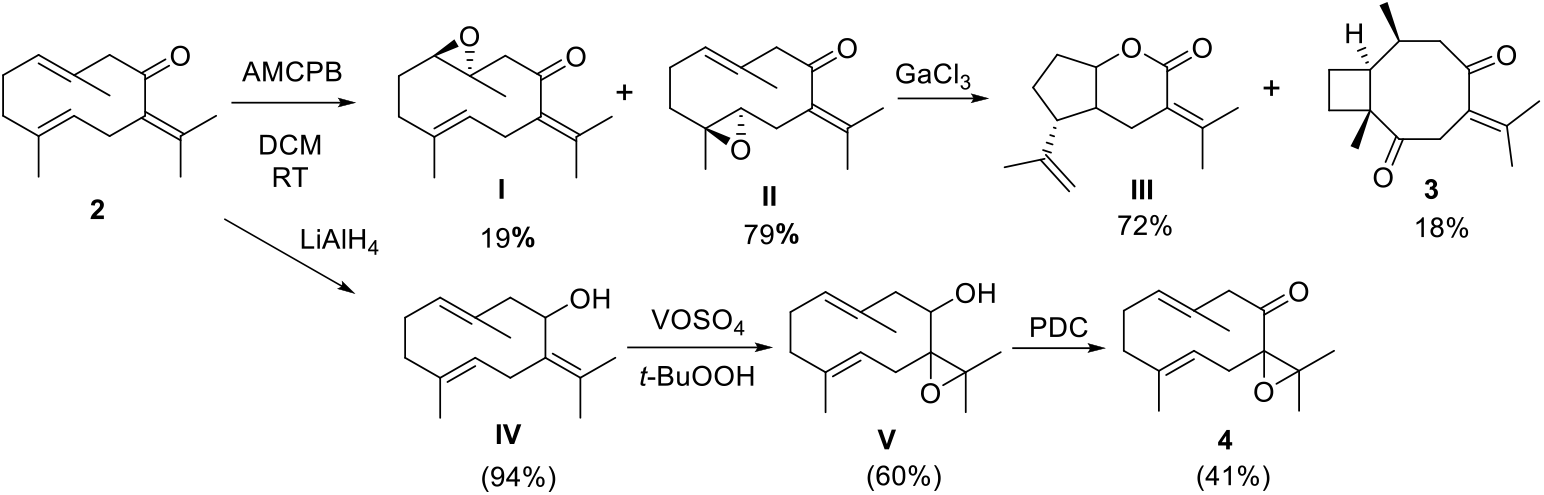
Synthesis of compounds **3** and **4** from germacrone.

Diketone (**3**) was produced after Ga(III)-mediated tandem cyclization-rearrangement of the 4-5-epoxide of germacrone, whereas the generation of **4** involved the reduction of germacrone with LAH followed by the hydroxyl-directed epoxidation of **IV** to form epoxide **V**, which was oxidized with PDC to produce **4**.

### 2.5. Evaluation of antiparasitic effects of G. macrorrhizum extracts and main components on extracellular protozoa

Anti-*Giardia* (AG) activity assays were conducted on *G. duodenalis* trophozoites from the ATCC® 30957 strain (assemblage A), using flat-bottom microwell plates in quadruplicate. Each well contained 150 μl of *G. duodenalis* trophozoites suspended in modified TYI-S-33 medium (Keister, 1983), supplemented with 10% heat-inactivated fetal bovine serum (FBS) (Sigma, Madrid, Spain) at a concentration of 10^6^ trophozoites/mL. Incubation was carried out for 24 hours at 37°C.

The anti-*Trichomonas* (AT) activity assay utilized *T. gallinae* trophozoites from an isolate obtained from a wood pigeon by Dr. M^a^ Teresa Gómez, employing round-bottom microwell plates in quadruplicate. Each well was filled with 150 μl of *T. gallinae* trophozoites in Trypticase-Yeast Extract-Maltose (TYM) medium, supplemented with 10% FBS at a concentration of 500,000 trophozoites/mL, with incubation for 24 hours at 37°C.

Anti-*Leishmania* (AL) activity was assessed on *L. infantum* promastigotes (MCAN/ES/98/LLM-722), kindly provided by Dr. J.M. Requena from CBM-CSIC, as well as on *L. infantum* (M/CAN/ES/96/BCN150 zymodeme MON-1), strain isolated from a dog with active visceral leishmaniasis (Fernández-Cotrina et al., 2013). *L. infantum* promastigotes (MCAN/ES/98/LLM-722) were cultured in RPMI medium supplemented with 15% FBS and 10 μg/mL of hemin (Acros Organics, Madrid, Spain), grown at 26°C. Promastigotes in logarithmic growth were seeded in 96-well flat-bottom plates at a volume of 100 μl of culture per well and incubated for 48 hours. Promastigotes for macrophage infection (M/CAN/ES/96/BCN150 zymodeme MON-1) were maintained at 26°C in Schneider’s insect medium (Lonza, Basel, Switzerland) supplemented with 10% FBS and 100 U/mL penicillin and 100 μg/mL streptomycin (Lonza, Basel, Switzerland) (Pen-Strep). Promastigotes in the stationary phase of growth (5–6 days) were washed twice in phosphate-buffered saline at 2,700 × g for 10 min at 22°C and employed for macrophage infection.

EOs and pure compounds were solved in DMSO (Sigma, Madrid, Spain) (<1% final concentration). The EOs and extracts were tested under concentrations of 800, 400, and 200 μg/mL, and in those cases that showed moderate antiparasitic activity (>50%) at the lowest concentration, additional testing was performed at lower concentrations: 100, and 50 μg/mL. Trophozoites/promastigotes viability was assessed using the modified MTT colorimetric assay, as described previously in the literature (Arroyo Díaz et al., 2023; Bailén et al., 2022, 2023), with results further validated by microscopic examination. The plate was analyzed in a spectrophotometer at a wavelength of 570 nm. The percentage of antiparasitic activity was calculated as growth inhibition, calculated using the following formula: % AT = 100−[(Ap− Ab)÷(Ac − Ab)]×100, where Ap is the absorbance of the tested product, Ab the absorbance of the blank and Ac the absorbance of the control wells (culture without treatment).

Pure compounds were tested using the same method, although at concentrations of 100, 75, 50, 25, 10, and 1 μg/mL. Metronidazole (Acros Organics, Madrid, Spain) served as a reference compound for anti-*Giardia* activity and Anti-*Trichomonas* activity and amphotericin B for antileishmanial activity.

### 2.6. Evaluation of antileishmanial effects of Germacrone on intracellular amastigotes

#### 2.6.1. Macrophages

Canine DH82 (ATCC®-CRL-10389TM) macrophages were maintained in Dulbecco’s modified Eagle’s medium (DMEM) (Thermo Fisher Scientific, Waltham, MA, USA) supplemented with 10% heat-inactivated FBS (Thermo Fisher Scientific), and Pen-Strep at 37 °C in 5% CO2.

#### 2.6.2. Cytotoxicity of germacrone on mammalian cells

DH82 canine macrophages (5 × 10^4^ per well) were seeded in 96-well plates in DMEM medium and kept overnight at 37 °C and 5% CO_2_. After washing with PBS, germacrone was added in varying concentrations (10, 50 and 100 μg/mL). After 48 h of treatment, cell viability was determined by the MTT method using a spectrophotometer (Multiskan EX, Thermo Electron Corporation, Finland) at 570 nm. The results were expressed as the cytotoxic concentration for 50% of the cell population (CC_50_), calculated using GraphPad Prism software version 8.30. Amphotericin B (GIBCO) at 1 μg/mL was used as the reference drug. The assays were performed in two independent experiments, each in quadruplicate.

#### 2.6.3. Infection index of macrophages

DH82 cells (5 × 10^4^ per well) were seeded in LabTek culture chamber slides (Thermo Scientific) and incubated overnight at 37 °C and 5% CO2. On the following day, macrophages were infected with *L. infantum* stationary phase-promastigotes at a ratio of 5:1 parasites/macrophage and after 4 h of incubation at 37°C, a time point that reflects initial infection, extracellular promastigotes were removed by washing. Germacrone was added at varying concentrations (10, 50 and 100 μg/mL) and macrophages were incubated in fresh medium for 48 h at 37°C and 5% CO2. After Giemsa staining, cells were mounted with Vitro-Clud (Deltalab SLU, Barcelona, Spain), and 400 macrophages were counted in duplicate in a microscope Olympus BX41. The percentage of infected macrophages and the mean of the number of amastigotes per infected macrophage (defined as the intensity of infection) were evaluated. The infection index was calculated by multiplication of both parameters to account for the overall parasite load (Domínguez-Bernal et al., 2014).

### 2.7. Cytotoxicity of pure compounds

African green monkey kidney cells (Vero cells) were maintained in Dulbecco’s modified Eagle’s minimal essential medium (DMEM) supplemented with 10% FBS and 1% Pen-Strep at 37°C under a humidified atmosphere of 5% CO2/95% air. Cells seeded in 96-well flat-bottom microplates with 100 μL medium per well (initial densities 10^4^ cells per well) were exposed for 48 h to serial dilutions (100, 75, 50, 25, 10 and 1 µg/mL) of the tested compounds in DMSO (<1% final concentration). Cell viability was analyzed by the MTT colorimetric assay method, and the purple-colored formazan precipitate was dissolved with 100 μL of DMSO (Bailén et al., 2022).

### 2.8. Morphological evaluation of the anti-protozoan effects

*L. infantum, T. gallinae* and *G. duodenalis* were studied at 0, 24, and 48 hours after treatment with germacrone (100 µg/mL) and controls (metronidazole at 10 µg/mL for *T. gallinae* and *G. duodenalis* and amphotericin B at 1 µg/mL for *L. infantum*).

For *L. infantum* and *T. gallinae*, cultures in the exponential growth phase (10^7^ promastigotes/mL and 5×10^5^ trophozoites/mL, respectively) were treated in 96-well microplates. At each time point, quadruplicates were assigned per condition: growth control, germacrone, and the corresponding positive control (metronidazole or amphotericin B). One sample per condition were centrifuged, resuspended in FBS and spread onto microscope slides. Once dried, samples were fixed and stained with Diff-Quik (Medion Diagnostic AG) following the manufacturer’s instructions. Additionally, three wells per condition (triplicates) were used for the MTT assay, with absorbance measured by spectrophotometry after 48 hours.

For *G. duodenalis*, Lab-Tek chamber slides (8 wells) were prepared with Keister culture medium and inoculated with trophozoites (10^6^ cells/mL). Each well contained 435 µl of culture medium and 15 µl of the test or control compounds at the above-mentioned concentrations. The chamber included: two wells for growth control (0.5 h, 48 h), two for germacrone (0.5 h, 48 h), and two for metronidazole (0.5 h, 48 h). Incubation was performed at 37°C. At each time point, the medium was removed, wells were dried, fixed, and stained with Diff-Quik.

Morphological changes were analyzed using light microscopy. For morphometric analysis, 30 randomly selected trophozoites per condition (treatment and controls at different times) were photographed using a researcher’s mobile phone through a calibrated microscope micrometer. Cell length, cell width, cell surface, nuclear length, nuclear width, axoneme (only *Giardia*), and length of axostyle (only *T. gallinae*) were measured with ImageJ software, and the data were recorded in an Excel database for statistical analysis.

### 2.9. Statistical analysis

Data was analyzed using STATGRAPHICS Centurion XIX (https://www.statgraphics.com). The mammalian cells and trophozoites viability were tested with each compound in a dose-response experiment to calculate their relative potency (CC_50_ or IC_50_ value, respectively). IC_50_ (μg/mL) expresses the dose of EOs, extracts or pure compounds needed to produce 50% mortality of trophozoites, while CC_50_ (μg/mL) expresses the dose of compounds necessary to produce 50% mortality of Vero cells.

Selectivity index was calculated for the antiparasitic activity of pure compounds, using the formula SI = IC_50_/CC_50_. Compounds with SI higher than one were considered as potential anti*-* parasitic compounds, since they are more toxic for protozoans than for mammalian cells.

Shapiro–Wilk test was used to assess the normality of the morphological variables. When data followed a normal distribution, the Student’s t-test was applied; otherwise, non-parametric tests, such as the Mann–Whitney U test, was used. Values of p< 0.05 were considered significant. For the morphological analysis, the mean, standard deviation, median, and interquartile range were calculated using SPSS version 20.

## 3. Results

### 3.1. Antiparasitic activity of extracts and essential oils from G. macrorrhizum

Six extracts and two EOs from *G. macrorrhizum* were tested on *T. gallinae* and *G. duodenalis* trophozoites, and *L. infantum* promastigotes, to evaluate their antiparasitic activity (Figure 2, Table 2).

**Table 2.**
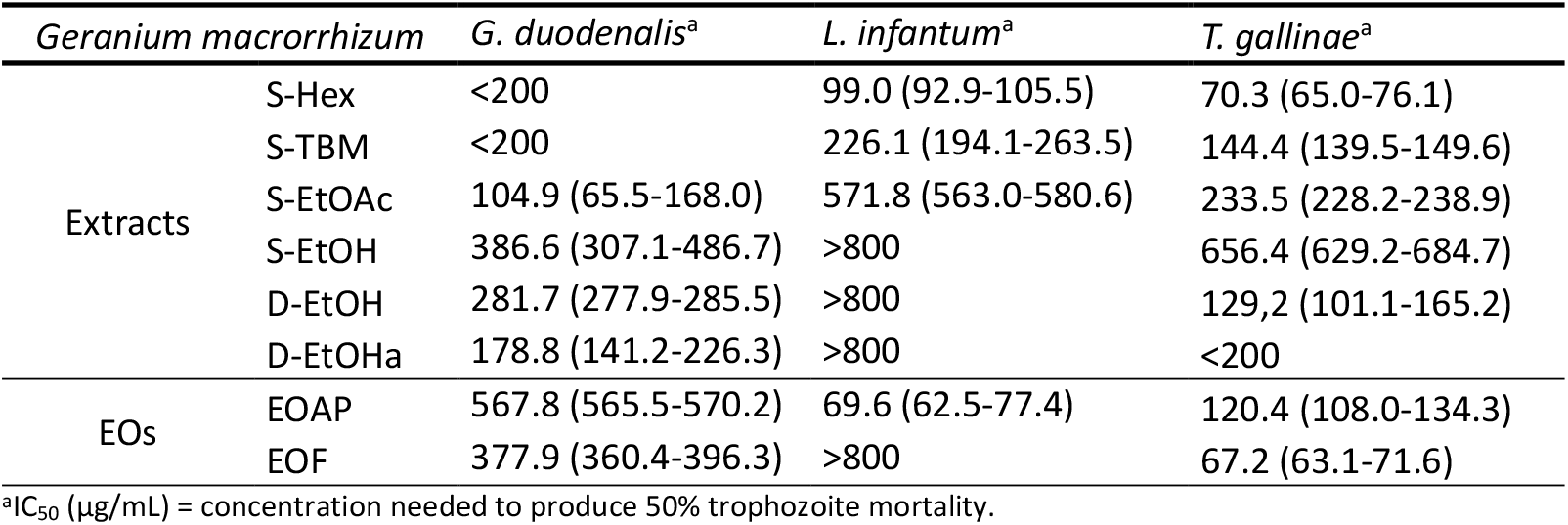
Effects of extracts and EOs from *G. macrorrhizum* on *L. infantum* promastigotes and *G. duodenalis* and *T. gallinae* trophozoites (IC_50_).

**Figure 2.**
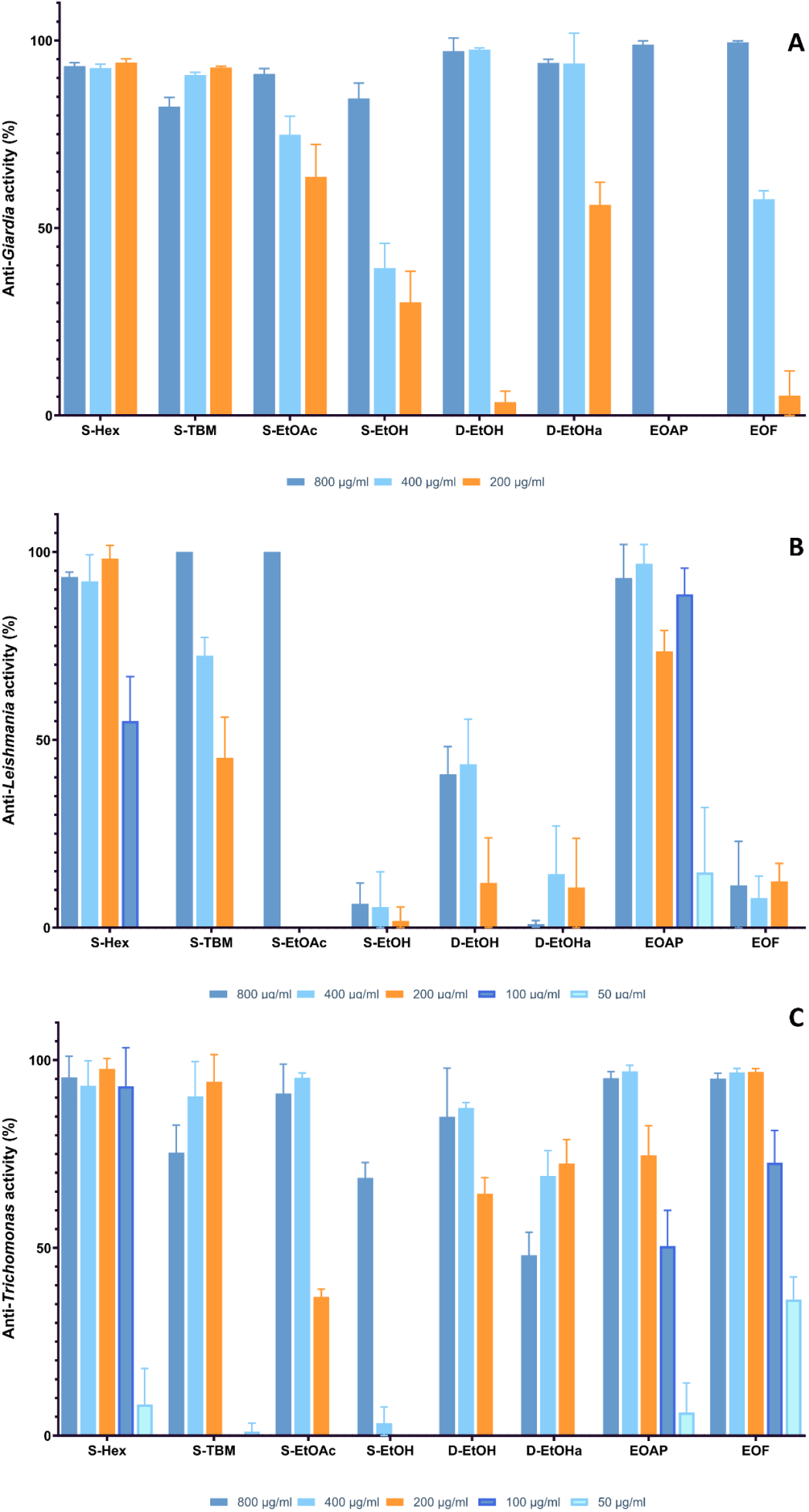
Percentage of anti-*Giardia* (A), anti-*Leishmania* (B) and anti-*Trichomonas* (C) activity of EOs and extracts from *G. macrorrhizum*.

Six extracts and two EOs exhibited activity against *G. duodenalis*, with efficacy increasing as solvent polarity decreases during the extraction process. Among the three ethanol-derived extracts (S-EtOH, D-EtOH, and D-EtOHa), those obtained by direct extraction (D-EtOH and D-EtOHa) exhibited stronger anti-*Giardia* activity. Notably, EOs from flowers demonstrated greater activity than those from aerial parts. Among all tested extracts, S-Hex was the most potent against *G. duodenalis* (IC_50_ = 90 μg/mL).

Similarly, against *L. infantum*, only three extracts and one essential oil (EOAP) displayed activity, again following an inverse relationship with polarity. No antileishmanial effects were observed in ethanol-extracted samples. The most effective compound in this case was the EO from aerial parts (EOAP) (IC_50_ = 69.6 μg/mL).

In the case of *T. gallinae*, six extracts and two EOs were active, making it the most sensitive parasite among those tested. As observed with the other protozoa, activity increased as polarity decreased, with the EO from flowers (EOF) exhibiting the highest efficacy (IC_50_ = 67.2 μg/mL).

Overall, S-Hex showed the strongest antiparasitic activity across all three protozoa, followed by S-TBM. Among the EOs, EOAP was the most effective against *L. infantum*, while EOF demonstrated the highest activity against *T. gallinae*.

### 3.2. Chemical composition of essential oils (EOs) and extracts from G. macrorrhizum

The chemical composition of two EOs from *G. macrorrhizum* aerial parts and flowers cultivated in greenhouse was determined by GC-MS (Table 3). Also, germacrone and β- elemenone were determined in the extracts obtained through sequential extraction.

**Table 3.** Chemical composition of extracts obtained through sequential extraction of EOs from *G. macrorrhizum* cultivated in a greenhouse.

| Extract/EO |  | Compounds (% Relative Abundance) |
| --- | --- | --- |
| EOs | EOAP | γ-elemene (3.05%); 1,2,2-trimethyl-1-(p-tolyl)-cyclopentane (3.72%); β-elemenone (**1**) (32.64%); nerolidol acetate (3.74%); germacrone (**2**) (40.72%) |
| | EOF | γ-elemene (1.11%); β-elemenone (**1**) (42.07%); nerolidol acetate (3.23%); germacrone (**2**) (36.85%) |
| Extracts | S-Hex | γ-elemene (0.86%); germacrone (**2**) (3.25%); 4,5 epoxygermacrone (16.00%) |
|  | S-TBM | germacrone (**2**) (1.31%) |
|  | S-EtOAc | - |
|  | S-EtOH | - |

β-Elemenone and germacrone were the main constituents of the analyzed essential oils (EOF and EOAP). β-Elemenone was more abundant in flowers, while germacrone predominated in aerial parts. Among the tested extracts, β-elemenone was not detected and germacrone was present in both S-Hex and S-TBM, with a higher concentration in S-Hex.

### 3.3. Antiparasitic and cytotoxic properties of G. macrorrhizum main compounds against extracellular protozoa

The main compounds of the S-TBM extract, elemenone (**1**) and germacrone (**2**) and germacrone derived-compounds, **3** and **4**, were also tested (Table 4). Among them, germacrone was effective against all three protozoa (*L. infantum, G. duodenalis*, and *T. gallinae*), whereas β- elemenone showed no activity against any.

**Table 4.** Antiparasitic effects of isolated compounds of *G. macrorrhizum* (IC_50_ (µg/mL)

| Compound | Vero cells | *L. infantum* (pm) |  | *G. duodenalis* |  | *T. gallinae* |  |
| --- | --- | --- | --- | --- | --- | --- | --- |
|  | CC<sub>50</sub> (μg/mL)<sup>a</sup> | IC<sub>50</sub> (μg/mL)<sup>b</sup> | SI<sup>c</sup> | IC<sub>50</sub> (μg/mL)<sup>b</sup> | SI<sup>c</sup> | IC<sub>50</sub> (μg/mL)<sup>b</sup> | SI<sup>c</sup> |
| **1** | >100 | >100 | 1.0 | >100 | 1 | >100 | 1.0 |
| **2** | ≈100 | 37.5 (33.6-41.9) | 2.7 | 55.2 (48.2-63.2) | 1.8 | 45.8 (37.2-56.3) | 2.2 |
| **3** | >100 | 35.5 (31.2-40.4) | 2.8 | >100 | 1 | >100 | 1.0 |
| **4** | >100 | 88.8 (62.6-125.9) | 1.1 | >100 | 1 | 51.6 (48.6-54.8) | 1.9 |
| Amphotericin B | 7.11 (Nasr et al., 2023) | 0.01 (Leal et al., 2013) | 711 | - | - | - | - |
| Metronidazole | >100 | - | - | 4.4 (3.5-5.5) | 22.7 | 1.0 (0.8-1.1) (Bailén et al., 2022) | 100 |

<sup>a</sup>CC<sub>50</sub> (μg/mL) = concentration needed to produce 50% Vero cell mortality; <sup>b</sup>IC<sub>50</sub> (μg/mL) = concentration needed to produce 50% trophozoite mortality. <sup>c</sup>SI: Selectivity index (CC<sub>50</sub>/IC<sub>50</sub>). pm = promastigotes.

Compound **3** exhibited activity exclusively against *L. infantum*, while **4** was effective against *T. gallinae* and displayed moderate activity against *L. infantum*. Overall, *L. infantum* was the most sensitive parasite and germacrone (**2**) the compound with the broader spectrum of action (Figure 3).

**Figure 3.**
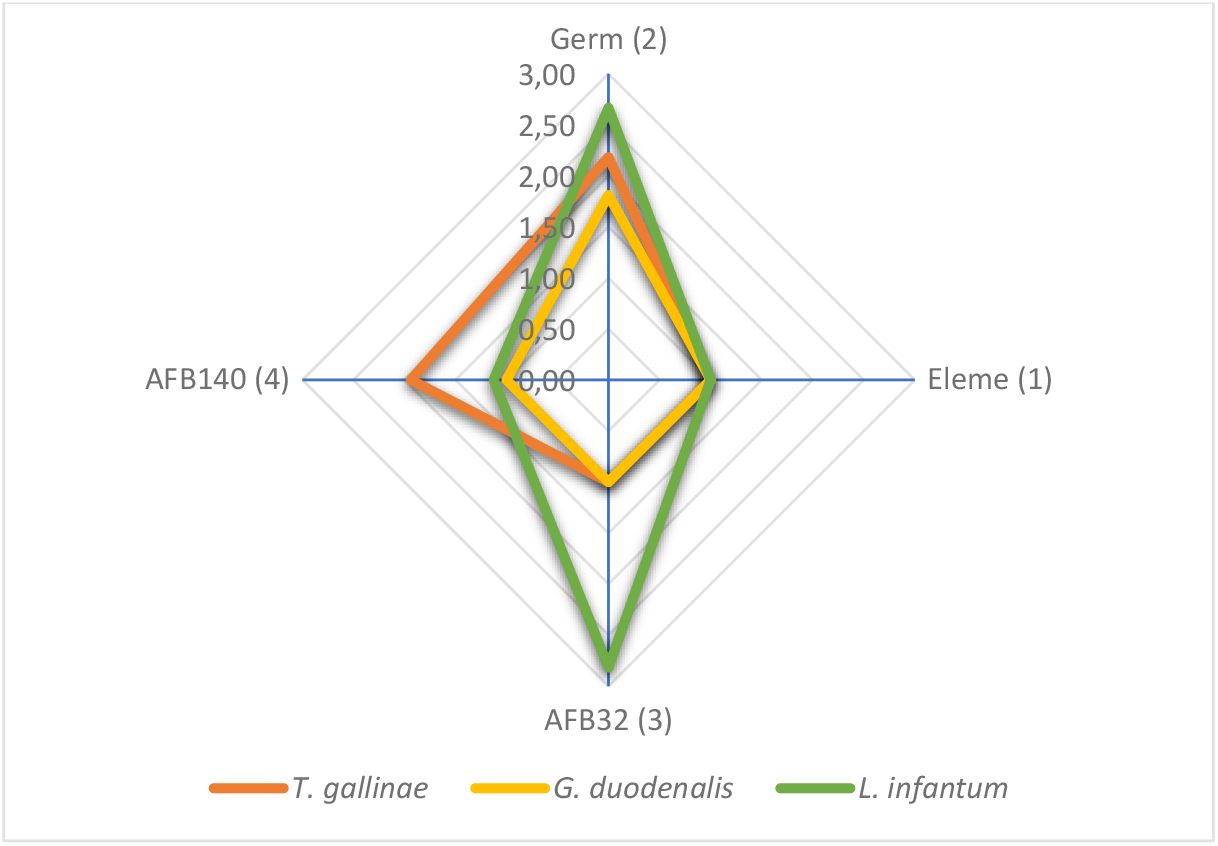
Selectivity indexes of main components of. *G. macrorrhizum* on extracellular protozoa (*T. gallinae* and *G. duodenalis* trophozoites and *L. infantum* promastigotes).

### 3.4. Antileishmanial effects of germacrone on intracellular amastigotes

The antileishmanial effects of germacrone were also evaluated on intracellular amastigotes in DH82 canine macrophages (Table 5). Prior to this, the cytotoxicity of germacrone was assessed in DH82 macrophages, yielding a CC_50_ value of 12.40 ± 1.03 μg/mL (Figure 4). At concentrations of 50 and 100 μg/mL, germacrone was lethal to canine macrophages.

**Table 5.** Antileishmanial effects of germacrone on intracellular amastigotes.

|  | % infected cells | n°amastigotes/<br>infected cell | infection index |
| --- | --- | --- | --- |
| control of infection | 42.8 ± 6.8 | 3.5 ± 1.8 | 152.1 ± 29.4 |
| DMSO 2% | 46.9 ± 6.9 | 2.7 ± 1.4 | 126.6 ± 4.9 |
| AmB (1 $\mu\text{g/mL}$) | 6.4 ± 3.7* | 1.4 ± 0.7* | 8.9 ± 0.0* |
| Germacrone (**2**) 10 $\mu\text{g/mL}$ | 25.9 ± 8.3* | 2.4 ± 1.1 | 61.0 ± 9.4* |
| Germacrone (**2**) 50 $\mu\text{g/mL}$ | dead cells | dead cells | dead cells |
| Germacrone (**2**) 100 $\mu\text{g/mL}$ | dead cells | dead cells | dead cells |

Data (mean ± SD) from two independent experiments are shown. (\* $p < 0.05$ ).

**Figure 4.**
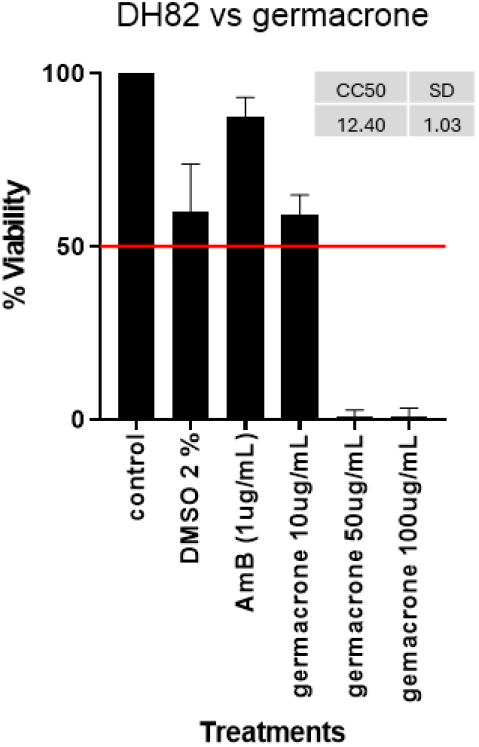
Cytotoxic effects of germacrone in DH82 canine macrophages (AmB: amphotericin B).

When evaluating the effect of germacrone on amastigote-infected canine macrophages, the percentage of infected cells decreased from 42.8% (control) to 25.9%, resulting in a reduction of the infection index by more than 50% (from 152.1 to 61.0).

### 3.5. Morphological evaluation of the anti-protozoan effects of germacrone

The antiprotozoal effects of germacrone on *G. duodenalis* and *T. gallinae* trophozoites and *Leishmania infantum* promastigotes at 48 h of exposition were further analyzed by light microscopy to detect morphological changes (Figure 5). In *G. duodenalis* trophozoites, germacrone provoked a less defined cytoplasm and nucleus of the cells. In *T. gallinae* trophozoites, an elongation of the cytoplasm and the nucleus could be observed. The effect on *L. infantum* promastigotes was less pronounced.

**Figure 5.**
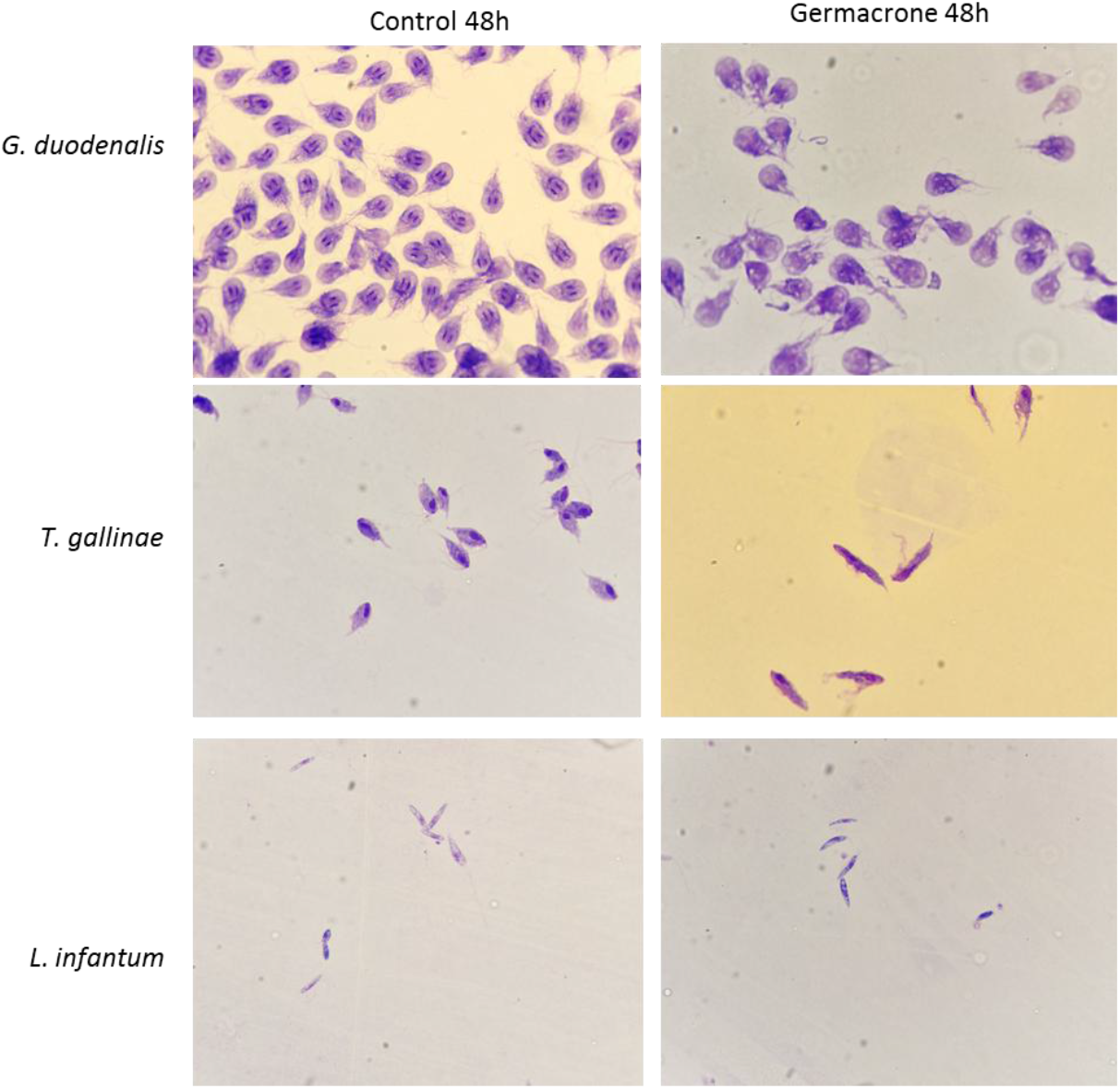
Effect of germacrone on *G. duodenalis* and *T. gallinae* trophozoites and *L. infantum* promastigotes stained with Diff-Quik after 48 h of exposition observed under light microscopy (x100).

#### 3.5.1. Giardia duodenalis

The effects of germacrone and metronidazole on various cellular parameters on *G. duodenalis* were analyzed throughout the study (Figure 6). For cell length, both germacrone and metronidazole showed significant reductions at 30 minutes of exposure compared to the control group. Regarding cell width, no differences were observed at 0.5 h with germacrone, but after 48 hours, a significant increase in cell width was noted. In contrast, metronidazole led to a decrease in cell width at 0.5 h when compared to the control. For nuclear length, both the left and right nuclear lengths decreased significantly with germacrone at 0.5 h and later increased after 48 h, relative to the control. No effects were observed for metronidazole on nuclear length. Regarding nuclear width, both germacrone and metronidazole caused a reduction at 0.5 h, followed by an increase at 48 h when compared to the control. The axoneme length decreased at 0.5 h for both germacrone and metronidazole; however, at 48 h, germacrone caused an increase, while metronidazole continued to decrease compared to the control. Finally, only metronidazole induced a reduction in cell surface area relative to the control within 0.5 h of exposure.

**Figure 6.**
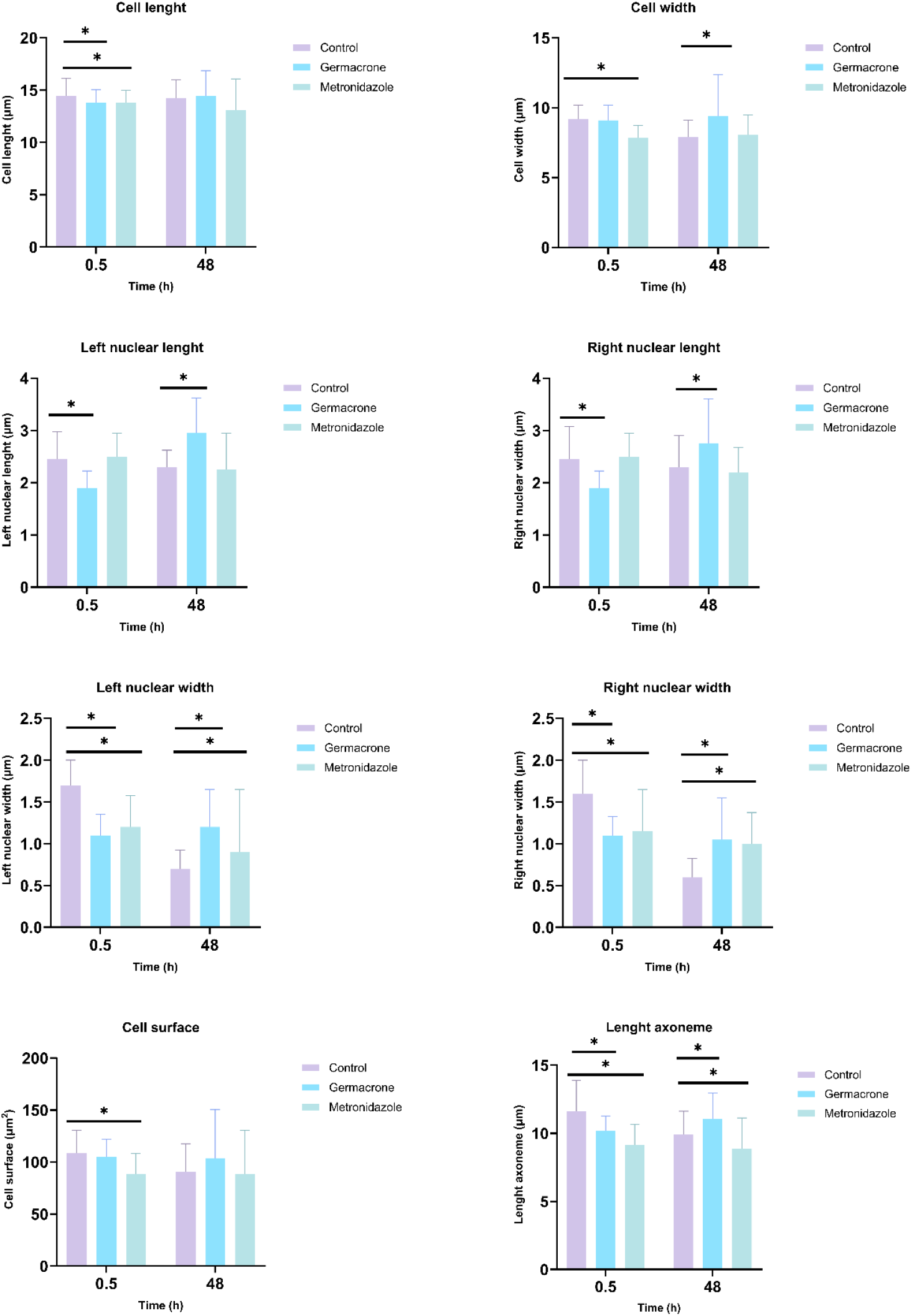
Effect of germacrone and metronidazole on *G. duodenalis* morphology (cell length, cell width, cell surface, length of axoneme, nuclear length and nuclear width) after 0.5 and 48 hours of exposure.

#### 3.5.2. Trichomonas gallinae

Optical microscopy analysis of *T. gallinae* trophozoites revealed significant morphological alterations following treatment with germacrone and metronidazole (Figure 7). Regarding cell length, germacrone induced an increase after 48 h, whereas metronidazole caused an initial increase at 0.5 h, followed by a reduction at 48 h. Cell width exhibited a significant decrease exclusively in germacrone-treated parasites compared to the control at 0.5 h of exposure. Nuclear length increased upon germacrone exposure but decreased with metronidazole, whereas nuclear width diminished at 0.5 h for both compounds. By 48 h, nuclear width increased in germacrone-treated trophozoites but further decreased in those exposed to metronidazole. Cell surface area decreased transiently at 0.5 h but increased at 48 h with germacrone, whereas metronidazole induced a significant reduction only at 48 h. Axostyle length decreased at 0.5 h for both treatments; however, after 48 h, it increased in germacrone-exposed parasites. Notably, the axostyle became undetectable in most metronidazole-treated trophozoites by 48 h.

**Figure 7.**
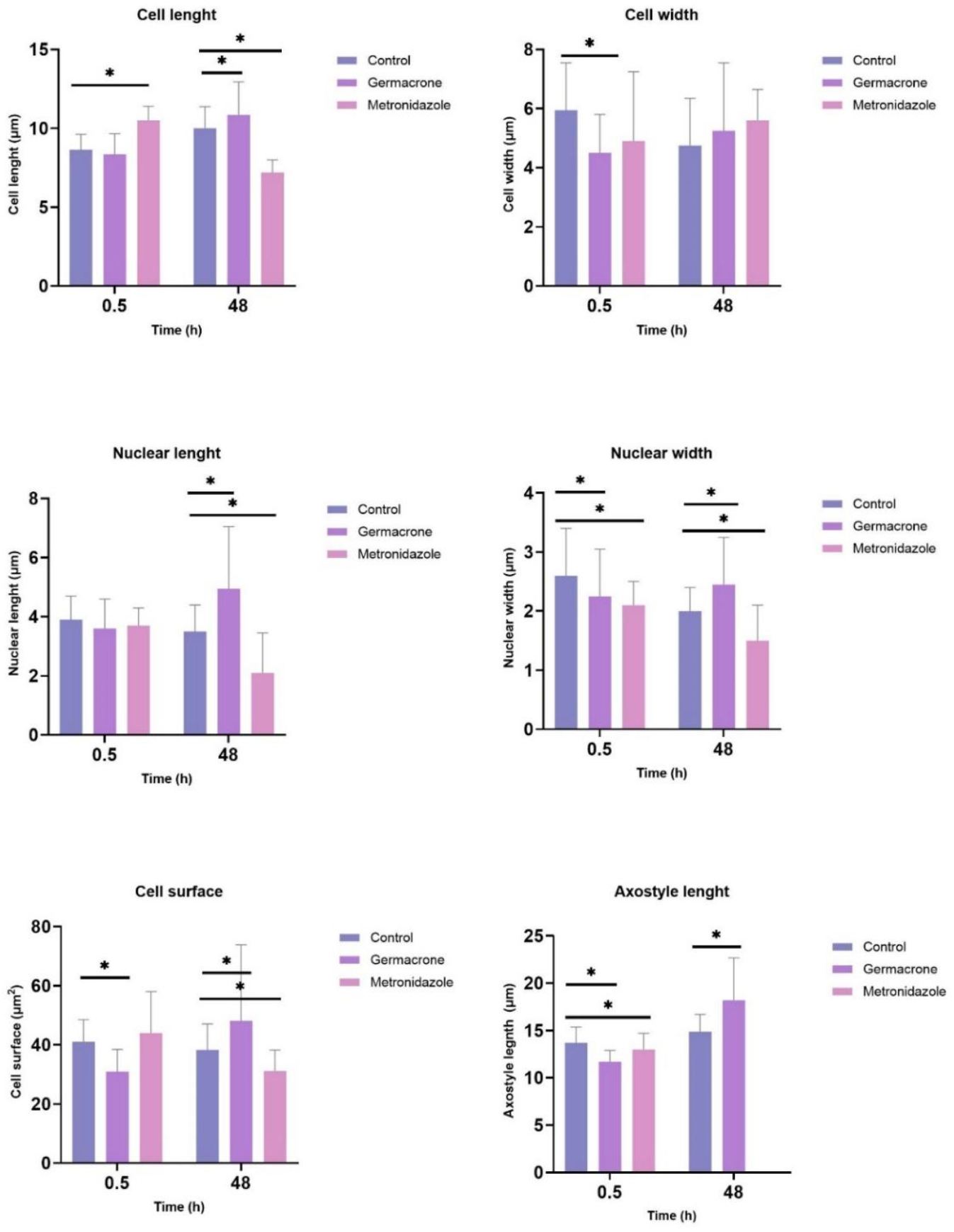
Effect of germacrone and metronidazole on *Trichomonas gallinae* morphology (cell length, cell width, cell surface, length of axostyle, nuclear length and nuclear width) after 0.5 and 48 hours of exposure.

**Figure 8.**
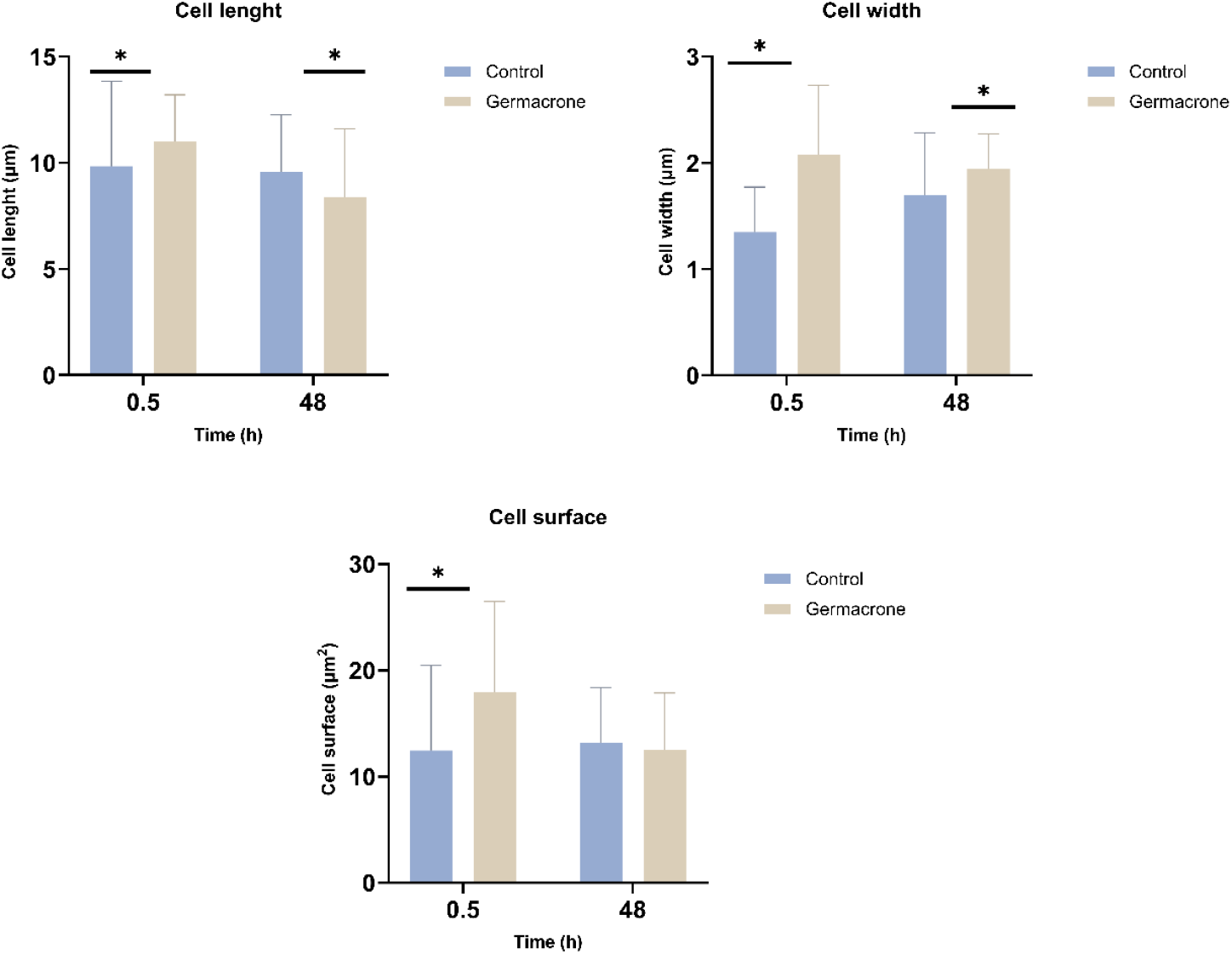
Effect of germacrone on *Leishmania infantum* promastigotes morphology (cell length, cell width and cell surface after 0.5 and 48 hours of exposure.

#### 3.5.3. Leishmania infantum

Optical microscopy analysis of *L. infantum* promastigotes revealed time-dependent morphological changes following germacrone exposure at 0.5 and 48 h. Cell length exhibited an initial increase at 0.5 hours but subsequently decreased at 48 h compared to the control. In contrast, cell width increased significantly at both 0.5 and 48 h post-treatment. Cell surface area transiently increased at 0.5 hours with no significant changes detected at 48 hours relative to the control. Amphotericin B treated promastigotes suffered strong morphological alterations that impede taking measures of the structures.

## 4. Discussion

The genus *Geranium* L. (eng. Cranesbills) comprises approximately 250 species of herbaceous plants that inhabits the Norther Hemisphere and tropical mountainous regions (Ilić et al., 2020). Despite demonstrating antimicrobial, insecticidal, ixodicidal, antioxidant, hypotensive, and sedative properties, the antiprotozoal potential of *Geranium macrorrhizum* extracts and EOs remains uninvestigated (Miliauskas et al., 2004; Navarro-Rocha et al., 2018; Radulović et al., 2012; Tzanova et al., 2024).

In the present study, the evaluation of extracts and EOs from *G. macrorrhizum* revealed broad-spectrum activity against *T. gallinae, G. duodenalis*, and *L. infantum*. Antiparasitic efficacy correlated inversely with solvent polarity, with non-polar extracts and EOs exhibiting the highest potency, suggesting that lipophilic constituents are likely responsible for the observed bioactivity. Ethanol extracts obtained via direct extraction (D-EtOH, D-EtOHa) showed greater efficacy compared to those obtained through sequential extraction (S-EtOH), potentially due to the prior depletion of active lipophilic compounds by non-polar solvents. Among the tested protozoa, *T. gallinae* was the most susceptible, with extract S-Hex demonstrating the highest efficacy. These findings are consistent with previous reports of antiprotozoal activity in related *Geranium* species, supporting the ethnopharmacological relevance of this genus. Methanolic root extracts of *Geranium niveum* have demonstrated activity against *G. duodenalis* (IC_50_ = 20.64 µg/mL) and *Entamoeba histolytica* (IC_50_ = 8.70 µg/mL) (Calzada et al., 1998). Similarly, *Geranium mexicanum* methanolic extract exhibited activity against *Trichomonas vaginalis* (IC_50_ = 56.0 µg/mL) (Calzada et al., 2007), while its dichloromethane–methanol root extracts showed moderate efficacy against *G. duodenalis* (IC_50_ = 100.4 µg/mL) (Calzada et al., 2005). In addition, ethanolic and methanolic extracts of *Geranium thunbergii* have shown potent antimalarial activity (Jun et al., 2025), and leaf extracts of *Geranium wallichianum* have demonstrated antileishmanial effects against both amastigote and promastigote forms of *Leishmania tropica* (Abbasi et al., 2019). Collectively, these studies highlight the broad antiparasitic potential of *Geranium* species and support further investigation into their bioactive constituents.

β-Elemenone and germacrone were identified as the major constituents of the analyzed EOs from aerial parts and flowers, with germacrone also detected in the less polar extracts S-Hex and S-TBM. Interestingly, although lipophilic extracts and EOs presented higher antiprotozoal activity compared to their more polar counterparts, the abundance of germacrone did not consistently correlate with antiparasitic efficacy, indicating that other compounds may contribute significantly to the observed effects.

Germacrone and elemenone, along with germacrone-derived compounds (compounds **3** and **4**), were evaluated for antiprotozoal activity. Significantly, none exhibited cytotoxicity against Vero cells (SI > 1), indicating selective toxicity toward protozoan parasites over mammalian cells. However, germacrone exhibited cytotoxicity against canine DH82 macrophages, which may be attributed to the use of an increased concentration of DMSO (2%) as a solvent, since DMSO alone showed similar levels of toxicity as germacrone. Germacrone demonstrated potent activity against *G. duodenalis* and *T. gallinae* trophozoites, as well as both promastigotes and amastigotes of *L. infantum*. Among the tested parasites, *L. infantum* was the most sensitive, with germacrone (1) emerging as the compound with the broadest spectrum of antiprotozoal activity. For this reason, germacrone was selected for further studies, such as the evaluation of morphological alterations.

Germacrone has emerged as a multifunctional natural compound with a broad spectrum of biological activities. It has shown notable anticancer potential by inducing cell cycle arrest and apoptosis in various malignancies, including breast, brain, liver, skin, prostate, gastric, and esophageal cancers, primarily through the modulation of critical signalling pathways and molecular targets involved in tumor progression (Riaz et al., 2020). Beyond its anticancer effects, germacrone exhibits antiviral activity, effectively inhibiting the replication of pseudorabies virus (He et al., 2019) and porcine reproductive and respiratory syndrome virus (Feng et al., 2016). Its insecticidal, acaricidal, and ixodicidal properties have also been documented (Galisteo Pretel et al., 2019; Navarro-Rocha et al., 2018) supporting its potential as a bioactive agent in pest control. Additionally, germacrone has been identified as an inhibitor of the hepatic cytochrome P450 isoform CYP2B6, suggesting possible implications for drug metabolism and pharmacokinetic interactions (Liu et al., 2024).

Germacrone is a naturally derived, biologically active sesquiterpene, which participates in a range of chemical transformations, including transannular cyclization, epoxidation processes, and photoinduced isomerization reactions (Barrero et al., 2008; Galisteo Pretel et al., 2019). The cyclization and further epoxidation of germacrone lead to compounds **3** and **4** modifying the antiprotozoal activity. The diketone **3** lost the anti-*Trichomonas* and anti-*Giardia* activity whilst compound **4** lost the anti-*Giardia* activity and presented a reduction in anti- *Leishmania* activity, which became moderate. Compounds **3** and **4** have previously been evaluated for their acaricidal and insecticidal properties. Compound **3** exhibited bioactivity comparable to that of germacrone in both insecticidal and acaricidal assays. In contrast, compound **4** demonstrated repellent activity against *Rhopalosiphum padi* and moderate acaricidal effects, although its overall efficacy was lower than that of germacrone (Galisteo Pretel et al., 2019).

Several EOs containing germacrone have demonstrated antiparasitic activity, highlighting the relevance of this sesquiterpene in natural product-based therapeutics. The EO of *Psidium brownianum*, which contains approximately 16% germacrone, exhibited significant activity against promastigotes of *L. braziliensis* and *L. infantum*, with IC_50_ values of 37.53 µg/mL and 75.83 µg/mL, respectively (Bezerra et al., 2022). Similarly, the EO of *Eugenia uniflora*, containing 8.52% germacrone, showed inhibitory effects on *L. amazonensis* promastigotes (da Silva et al., 2018). In addition to germacrone, other sesquiterpenes have been reported to possess antiparasitic properties. For instance, γ-elemene and curzerene have shown activity against both amastigote and promastigote forms of *L. amazonensis* (de Lima Nunes et al., 2021a). Other sesquiterpenes (euparin and santhemoidin C) or sequiterpene lactones (anhydroartemorin, cis,trans-costunolide-14-acetate, and 4-hydroxyarbusculin A) demonstrated efficacy against *L. infantum* (Elso et al., 2021), *L. amazonensis*, and *L. donovani* (Bethencourt-Estrella et al., 2021), respectively. The antiparasitic mechanisms of sesquiterpenes against *Leishmania* spp. are thought to involve the induction of apoptosis, necrosis, autophagy, and disruption of the parasite cell membrane, leading to metabolic and structural alterations that compromise parasite survival and proliferation (Rodrigues et al., 2023).

Beyond leishmanicidal activity, various sesquiterpenes have also demonstrated efficacy against other protozoan parasites. In the case of *Giardia* sp., antiparasitic effects have been reported for incomptine A analogs (Bautista et al., 2014), cadinane-type sesquiterpenes (Rodríguez-Chávez et al., 2015), β-caryophyllene-4,5-α-oxide (Calzada et al., 2001), brevilin A (Yu et al., 1994), and neurolenin-type furanoheliangolides such as neurolenin B (Passreiter et al., 1999). Regarding *T. vaginalis*, notable activity has been observed for a chloro derivative of α- santonin (Ivasenko et al., 2006), dihydroartemisinin (Tang et al., 2010), and caryophyllene oxide (Cheikh-Ali et al., 2011). These findings underscore the broad-spectrum antiparasitic potential of sesquiterpenes and their derivatives, supporting their continued exploration as candidates for novel antiparasitic agents.

Besides the activity of germacrone on the tested protozoa metabolism, the morphological effects were also evaluated. Germacrone exerted a strong effect on the morphology of the mentioned flagellates, and the activity across *L. infantum, G. duodenalis*, and *T. gallinae* reveal both common and species-specific responses. In all three species, germacrone induced time-dependent alterations in cell shape and size, suggesting a conserved impact on cytoskeletal dynamics. In *L. infantum*, germacrone caused an early increase in cell length and width, followed by a reduction in length at 48 h, indicating transient elongation and sustained cell broadening. This contrasts with *G. duodenalis*, where germacrone led to early shortening and delayed widening, and with *T. gallinae*, where cell length increased only at 48 h and width decreased early. *G. duodenalis* and *T. gallinae* also showed transient nuclear narrowing followed by recovery. Germacrone promoted axoneme/axostyle elongation in both *Giardia* and *Trichomonas* respectively, suggesting a potential stabilizing effect on microtubule-associated structures. In contrast, metronidazole consistently induced rapid and sustained morphological damage in both protozoa, including axostyle/axoneme loss and nuclear shrinkage. These findings suggest that germacrone elicits a more dynamic and potentially reversible response across different protozoan parasites, whereas metronidazole exerts a more immediate and degenerative effect. The differential responses observed highlight the importance of parasite-specific structural features in shaping drug sensitivity and may inform the development of broad-spectrum antiparasitic strategies.

The specific mode of action of germacrone against protozoa has not been detailed so far, but other natural terpenes with anti-*Giardia* activity have different mechanisms of action. Andrographolide, a diterpenoid lactone, caused significant alterations in *G. duodenalis* trophozoite shape and size and effectively inhibited trophozoite adhesion (Haldar et al., 2024). Other molecules, such as linearolactone, a neo-clerodane-type diterpene, induced partial arrest in the S phase of the trophozoite cell cycle without evidence of reactive oxygen species (ROS) production. It also triggered pronecrotic cell death and ultrastructural alterations, including changes in vacuole abundance, the appearance of perinuclear and periplasmic spaces, and glycogen granule deposition (Argüello-García et al., 2022). Additionally, other terpenes like the sesquiterpene lactone dihydroartemisinin with activity against related protozoa, such as *T. vaginalis*, it disrupts membrane systems, further illustrating the mechanistic diversity of this compound class (Tang et al., 2010).

Several sesquiterpene lactones have demonstrated antileishmanial activity through diverse mechanisms. Helenalin, mexicanin, and dehydroleucodine inhibited the in vitro growth of *Leishmania* promastigotes, with evidence of DNA fragmentation, and in the case of helenalin, pronounced vacuolization was also observed (Barrera et al., 2008). Similarly, (-)-α-bisabolol showed efficacy against intracellular amastigotes of *L. tropica*, inducing oxidative stress and mitochondrial-dependent apoptosis without compromising plasma membrane integrity (Corpas-López et al., 2016). In contrast, γ-elemene exhibited direct activity against both promastigote and amastigote forms of *Leishmania* (L.) *amazonensis*, primarily by disrupting plasma membrane integrity (de Lima Nunes et al., 2021b). These findings highlight the mechanistic diversity of sesquiterpene lactones and their potential as antiparasitic agents targeting different cellular pathways in *Leishmania*. Altogether, it seems that terpenes and sesquiterpenes exert a remarkable effect on both, the metabolic activity of the protozoa and the protozoa architecture, as we have observed with germacrone in the present study.

## 5. Conclusions

The results of this study demonstrate that extracts and EOs from *Geranium macrorrhizum*, particularly those rich in germacrone, exhibit promising antiparasitic activity against *G. duodenalis, T. gallinae*, and *L. infantum*. The inverse correlation between solvent polarity and antiparasitic efficacy highlights the relevance of lipophilic constituents, with germacrone emerging as a key bioactive compound. Its broad-spectrum activity, combined with selective toxicity toward protozoan parasites (SI > 1), supports its potential as a lead compound for therapeutic development. These findings place *G. macrorrhizum* as a valuable source of natural antiprotozoal agents and warrant continued investigation into its active constituents and mechanisms of action.

## 6. Conflict of Interest

The authors declare that the research was conducted in the absence of any commercial or financial relationships that could be construed as a potential conflict of interest.

## 7. Author Contributions

Conceptualization, AG, MTG and MB. Data curation, AG, MTG and MB. Formal analysis, SMH, MTG, MJI, JC, AG y MB. Funding acquisition, AG, and MTG. Investigation SMH, MJIG, IAC, RSL, JC and JA. Methodology, AFB, EO, JFQ, JNR, JC, AG, MTG and MB. Resources, AG, JNR, AFB, EO, JFQ, JC, MTG and MB. Supervision: AG, MB and MTG. Writing – original draft, MTG, MB. Writing, reviewing and editing, AG, MTG and MB.

## 8. Funding

This work has been partially financed by Grants PID2020-114207RB-I00 (Spanish Ministry of Science and Innovation), MINISTERIO DE CIENCIA, PID2019-106222RB-C32/SRA (State Research Agency, 10.13039/501100011033), PID2019-106222RB-C31/SRA (State Research Agency, 10.13039/501100011033, and Unidad Asociada UGR-CSIC BIOPLAG.

## 9. Acknowledgments

We want to thank Rubén Muñoz for his kind assistance in conducting identification of chemical components of essential oils.

## Notes

### Competing Interest Statement

The authors have declared no competing interest.

## References

Abbasi, B. A., Iqbal, J., Ahmad, R., Zia, L., Kanwal, S., Mahmood, T., Wang, C., & Chen, J. T. (2019). Bioactivities of Geranium wallichianum Leaf Extracts Conjugated with Zinc Oxide Nanoparticles. Biomolecules 2020, Vol. 10, Page 38, 10(1), 38. 10.3390/BIOM10010038

Aedo, C., Garmendia, F. M., & Pando, F. (1998). World checklist of Geranium L. (Geraniaceae). Anales Del Jardin Botanico de Madrid, 56(2), 211–252. 10.3989/ajbm.1998.v56.i2.230

Amin, A., Bilic, I., Liebhart, D., & Hess, M. (2014). Trichomonads in birds-a review. Parasitology, 141(6), 733–747. 10.1017/S0031182013002096

Argüello-García, R., Calzada, F., Chávez-Munguía, B., Matus-Meza, A. S., Bautista, E., Barbosa, E., Velazquez, C., Hernández-Caballero, M. E., Ordoñez-Razo, R. M., & Velázquez-Domínguez, J. A. (2022). Linearolactone Induces Necrotic-like Death in Giardia intestinalis Trophozoites: Prediction of a Likely Target. Pharmaceuticals, 15(7). 10.3390/PH15070809

Argüello-García, R., Leitsch, D., Skinner-Adams, T., & Ortega-Pierres, M. G. (2020). Drug resistance in Giardia: Mechanisms and alternative treatments for Giardiasis. Advances in Parasitology, 107, 201–282. 10.1016/bs.apar.2019.11.003

Arroyo Díaz, J. E., Gómez Muñoz, M. T., Martínez-Díaz, R. A., & González-Coloma, A. (2023). Adaptación del ensayo colorimétrico del MTT para la evaluación de la actividad frente a Giardia duodenalis. Revista Argentina de Parasitología, 12. http://www.revargparasitologia.com.ar/ojs/index.php/rap/article/view/4

Atanasov, A. G., Zotchev, S. B., Dirsch, V. M., Orhan, I. E., Banach, M., Rollinger, J. M., Barreca, D., Weckwerth, W., Bauer, R., Bayer, E. A., Majeed, M., Bishayee, A., Bochkov, V., Bonn, G. K., Braidy, N., Bucar, F., Cifuentes, A., D’Onofrio, G., Bodkin, M., … Supuran, C. T. (2021). Natural products in drug discovery: advances and opportunities. Nature Reviews Drug Discovery, 20(3), 200–216. 10.1038/S41573-020-00114-Z

Bailén, M., Díaz-Castellanos, I., Azami-Conesa, I., Alonso Fernández, S., Martínez-Díaz, R. A., Navarro-Rocha, J., Gómez-Muñoz, M. T., & González-Coloma, A. (2022). Anti-Trichomonas gallinae activity of essential oils and main compounds from Lamiaceae and Asteraceae plants. Frontiers in Veterinary Science, 9. 10.3389/FVETS.2022.981763

Bailén, M., Illescas, C., Quijada, M., Martínez-Díaz, R. A., Ochoa, E., Gómez-Muñoz, M. T., Navarro-Rocha, J., & González-Coloma, A. (2023). Anti-Trypanosomatidae Activity of Essential Oils and Their Main Components from Selected Medicinal Plants. Molecules, 28(3), 1467. 10.3390/MOLECULES28031467/S1

Barrera, P. A., Jimenez-Ortiz, V., Tonn, C., Giordano, O., Galanti, N., & Sosa, M. A. (2008). Natural Sesquiterpene Lactones Are Active against Leishmania mexicana. Source: The Journal of Parasitology, 94(5), 1143–1149. https://about.jstor.org/terms

Barrero, A. F., Mar Herrador, M., Arteaga, P., & Catalán, J. V. (2008). Germacrone: Occurrence, Synthesis, Chemical Transformations and Biological Properties. Natural Product Communications, 3(4), 567–576.

Bautista, E., Calzada, F., López-Huerta, F. A., Yépez-Mulia, L., & Ortega, A. (2014). Antiprotozoal activity of 8-acyl and 8-alkyl incomptine A analogs. Bioorganic & Medicinal Chemistry Letters, 24(15), 3260–3262. 10.1016/J.BMCL.2014.06.017

Bethencourt-Estrella, C. J., Nocchi, N., López-Arencibia, A., Nicolás-Hernández, D. S., Souto, M. L., Suárez-Gómez, B., Díaz-Marrero, A. R., Fernández, J. J., Lorenzo-Morales, J., & Piñero, J. E. (2021). Antikinetoplastid activity of sesquiterpenes isolated from the zoanthid Palythoa aff. Clavata. Pharmaceuticals, 14(11), 1095. 10.3390/PH14111095/S1

Bezerra, J. N., Gomez, M. C. V., Rolón, M., Coronel, C., Almeida-Bezerra, J. W., Fidelis, K. R., Menezes, S.A.de, Cruz R. P. da, Duarte, A. E., Ribeiro, P. R. V., Brito, E.S.de, Coutinho, H. D. M., Morais-Braga, M. F. B., & Bezerra, C. F. (2022). Chemical composition, Evaluation of Antiparasitary and Cytotoxic Activity of the essential oil of Psidium brownianum MART EX. DC. Biocatalysis and Agricultural Biotechnology, 39, 102247. 10.1016/J.BCAB.2021.102247

Calzada, F., Cedillo-Rivera, R., & Mata, R. (2001). Antiprotozoal activity of the constituents of Conyza filaginoides. Journal of Natural Products, 64(5), 671–673. 10.1021/NP000442O

Calzada, F., Cervantes-Martínez, J. A., & Yépez-Mulia, L. (2005). In vitro antiprotozoal activity from the roots of Geranium mexicanum and its constituents on Entamoeba histolytica and Giardia lamblia. Journal of Ethnopharmacology, 98(1–2), 191–193. 10.1016/j.jep.2005.01.019

Calzada, F., Meckes, M., Cedillo-Rivera, R., Tapia-Contreras, A., & Mata, R. (1998). Screening of Mexican medicinal plants for antiprotozoal activity. Taylor & FrancisF Calzada, M Meckes, R Cedillo-Rivera, A Tapia-Contreras, R MataPharmaceutical Biology, 1998•Taylor & Francis, 36(5), 305–309. 10.1076/PHBI.36.5.305.4653

Calzada, F., Yépez-Mulia, L., & Tapia-Contreras, A. (2007). Effect of Mexican medicinal plant used to treat trichomoniasis on Trichomonas vaginalis trophozoites. Journal of Ethnopharmacology, 113(2), 248–251. 10.1016/j.jep.2007.06.001

Cernikova, L., Faso, C., & Hehl, A. B. (2018). Five facts about Giardia lamblia. PLOS Pathogens, 14(9), e1007250. 10.1371/JOURNAL.PPAT.1007250

Cheikh-Ali, Z., Adiko, M., Bouttier, S., Bories, C., Okpekon, T., Poupon, E., & Champy, P. (2011). Composition, and antimicrobial and remarkable antiprotozoal activities of the essential oil of rhizomes of Aframomum sceptrum K. Schum. (Zingiberaceae). Chemistry and Biodiversity, 8(4), 658–667. 10.1002/CBDV.201000216

Corpas-López, V., Merino-Espinosa, G., Díaz-Sáez, V., Morillas-Márquez, F., Navarro-Moll, M. C., & Martín-Sánchez, J. (2016). The sesquiterpene (−)-α-bisabolol is active against the causative agents of Old World cutaneous leishmaniasis through the induction of mitochondrial-dependent apoptosis. Apoptosis, 21(10), 1071–1081. 10.1007/S10495-016-1282-X/FIGURES/5

da Silva, V. P., Alves, C. C. F., Miranda, M. L. D., Bretanha, L. C., Balleste, M. P., Micke, G. A., Silveira, E. V., Martins, C. H. G., Ambrosio, M. A. L. V., de Souza Silva, T., Tavares, D. C., Magalhães, L. G., Silva, F. G., & Egea, M. B. (2018). Chemical composition and in vitro leishmanicidal, antibacterial and cytotoxic activities of essential oils of the Myrtaceae family occurring in the Cerrado biome. Industrial Crops and Products, 123, 638–645. 10.1016/J.INDCROP.2018.07.033

de Lima Nunes, T.A., Costa, L. H., De Sousa, J. M. S., De Souza, V. M. R., Rodrigues, R. R. L., Val, M. da C. A., Pereira, A. C. T. da C., Ferreira, G. P., Da Silva, M. V., Da Costa, J. M. A. R., Véras, L. M. C., Diniz, R. C., & Rodrigues, K. A. da F. (2021a). Eugenia piauhiensis Vellaff. essential oil and γ-elemene its major constituent exhibit antileishmanial activity, promoting cell membrane damage and in vitro immunomodulation. Chemico-Biological Interactions, 339, 109429. 10.1016/J.CBI.2021.109429

de Lima Nunes, T.A. de L., Santos, M. M., de Oliveira, M. S., de Sousa, J. M. S., Rodrigues, R. R. L., Sousa, P. S. de A., de Araújo, A. R., Pereira, A. C. T. da C., Ferreira, G. P., Rocha, J. A., Rodrigues Junior, V., da Silva, M. V., & Rodrigues, K. A. da F. (2021b). Curzerene antileishmania activity: Effects on Leishmania amazonensis and possible action mechanisms. International Immunopharmacology, 100, 108130. 10.1016/J.INTIMP.2021.108130

Domínguez-Bernal, G., Jiménez, M., Molina, R., Ordóñez-Gutiérrez, L., Martínez-Rodrigo, A., Mas, A., Cutuli, M. T., & Carrión, J. (2014). Characterisation of the ex vivo virulence of Leishmania infantum isolates from Phlebotomus perniciosus from an outbreak of human leishmaniosis in Madrid, Spain. Parasites and Vectors, 7(1), 1–7. 10.1186/S13071-014-0499-1/FIGURES/4

Elso, O. G., Clavin, M., Hernandez, N., Sgarlata, T., Bach, H., Catalan, C. A. N., Aguilera, E., Alvarez, G., & Sülsen, V. P. (2021). Antiprotozoal Compounds from Urolepis hecatantha (Asteraceae). Evidence-Based Complementary and Alternative Medicine, 2021(1), 6622894. 10.1155/2021/6622894

European Commission. (1995). Commission Regulation (EC) No 1798/95 of 25 July 1995 amending Annex IV to Council Regulation (EEC) No 2377/90 laying down a Community procedure for the establishment of maximum residue limits of veterinary medicinal products in foodstuffs of animal origin. Official Journal of the European Communities, L174, 20–21.

Feng, J., Bai, X., Cui, T., Zhou, H., Chen, Y., Xie, J., Shi, Q., Wang, H., & Zhang, G. (2016). In Vitro Antiviral Activity of Germacrone Against Porcine Reproductive and Respiratory Syndrome Virus. Current Microbiology, 73(3), 317–323. 10.1007/S00284-016-1042-8

Fernández-Cotrina, J., Iniesta, V., Belinchón-Lorenzo, S., Muñoz-Madrid, R., Serrano, F., Parejo, J. C., Gómez-Gordo, L., Soto, M., Alonso, C., & Gómez-Nieto, L. C. (2013). Experimental model for reproduction of canine visceral leishmaniosis by Leishmania infantum. Veterinary Parasitology, 192(1–3), 118–128. 10.1016/J.VETPAR.2012.10.002

Fraga, B. M., Díaz, C. E., Bolaños, P., Bailén, M., Andrés, M. F., & González-Coloma, A. (2020). Alkane-, alkene-, alkyne-γ-lactones and ryanodane diterpenes from aeroponically grown Persea indica roots. Phytochemistry, 176, 112398. 10.1016/J.PHYTOCHEM.2020.112398

Galisteo Pretel, A., Pérez Del Pulgar, H., Guerrero de León, E., López-Pérez, J. L., Olmeda, A. S., Gonzalez-Coloma, A. F, Barrero, A., & Quílez Del Moral, J.F. (2019). Germacrone Derivatives as new Insecticidal and Acaricidal Compounds: A Structure-Activity Relationship. Molecules, 24(16), 2898. 10.3390/MOLECULES24162898

Gervazoni, L. F. O., Barcellos, G. B., Ferreira-Paes, T., & Almeida-Amaral, E. E. (2020). Use of Natural Products in Leishmaniasis Chemotherapy: An Overview. Frontiers in Chemistry, 8, 579891. 10.3389/FCHEM.2020.579891/REFERENCE

Haldar, T., Sardar, S. K., Ghosal, A., Prasad, A., Nakano, Y. S., Dutta, S., Nozaki, T., & Ganguly, S. (2024). Andrographolide induced cytotoxicity and cell cycle arrest in Giardia trophozoites. Experimental Parasitology, 262, 108773. 10.1016/J.EXPPARA.2024.108773

He, W., Zhai, X., Su, J., Ye, R., Zheng, Y., & Su, S. (2019). Antiviral Activity of Germacrone against Pseudorabies Virus in Vitro. Pathogens 2019, Vol. 8, Page 258, 8(4), 258. 10.3390/PATHOGENS8040258

Ilić, M. D., Marčetić, M. D., Zlatković, B. K., Lakušić, B. S., Kovačević, N. N., & Drobac, M. M. (2020). Chemical Composition of Volatiles of Eight Geranium L. Species from Vlasina Plateau (South Eastern Serbia). Chemistry & Biodiversity, 17(2), e1900544. 10.1002/CBDV.201900544

Ivasenko, S. A., Edil’baeva, T. T., Kulyyasov, A. T., Atazhanova, G. A., Drab, A. I., Turdybekov, K. M., Raldugin, V. A., & Adekenov, S. M. (2006). Structure and biological activity of α-santonin chloro-derivatives. Chemistry of Natural Compounds, 42(1), 36–40. 10.1007/S10600-006-0031-8/METRICS

Jun, H., Han, J. H., Hong, M., Fitriana, F., Syahada, J. H., Lee, W. J., Mazigo, E., Louis, J. M., Nguyen, V. T., Cha, S. H., Chun, W., Park, W. S., Lee, S. J., Na, S., Lee, S. U., Han, E. T., Kwon, T. H., & Han, J. H. (2025). Ellagic Acid from Geranium thunbergii and Antimalarial Activity of Korean Medicinal Plants. Molecules, 30(2), 359. 10.3390/MOLECULES30020359/S1

Keister, D. B. (1983). Axenic culture of Giardia lamblia in TYI-S-33 medium supplemented with bile. Transactions of The Royal Society of Tropical Medicine and Hygiene, 77(4), 487–488. 10.1016/0035-9203(83)90120-7

Leal, S. M., Pino, N., Stashenko, E. E., Martínez, J. R., & Escobar, P. (2013). Antiprotozoal activity of essential oils derived from Piper spp. grown in Colombia. Journal of Essential Oil Research, 25(6), 512–519. 10.1080/10412905.2013.820669;JOURNAL:JOURNAL:TJEO20;REQUESTEDJOURNAL:JOURNAL:TJEO20;WGROUP:STRING:PUBLICATION

Liu, S., Zhao, Y., Tang, X., Yang, J., Pan, C., Liu, C., Han, J., Li, C., Yi, Y., Li, Y., Cheng, J., Zhang, Y., Wang, L., Tian, J., Wang, Y., Wang, L., & Liang, A. (2024). In vitro inhibition of six active sesquiterpenoids in zedoary turmeric oil on human liver cytochrome P450 enzymes. Journal of Ethnopharmacology, 322, 117588. 10.1016/J.JEP.2023.117588

Menezes, S. A., & Tasca, T. (2023). Essential Oils and Terpenic Compounds as Potential Hits for Drugs against Amitochondriate Protists. Tropical Medicine and Infectious Disease 2023, Vol. 8, Page 37, 8(1), 37. 10.3390/TROPICALMED8010037

Miliauskas, G., Van Beek, T. A., Venskutonis, P. R., Linssen, J. P. H., & De Waard, P. (2004). Antioxidative activity of Geranium macrorrhizum. European Food Research and Technology, 218(3), 253–261. 10.1007/S00217-003-0836-7/TABLES/3

Nasr, I. S. A., Koko, W. S., Khan, T. A., Schobert, R., & Biersack, B. (2023). Antiparasitic Activity of Fluorophenyl-Substituted Pyrimido[1,2-a]benzimidazoles. Biomedicines, 11(1), 219. 10.3390/BIOMEDICINES11010219/S1

Navarro-Rocha, J., F. Barrero, A., Burillo, J., Olmeda, A. S., & González-Coloma, A. (2018). Valorization of essential oils from two populations (wild and commercial) of Geranium macrorrhizum L. Industrial Crops and Products, 116, 41–45. 10.1016/J.INDCROP.2018.02.046

Passreiter, C. M., Sandoval-Ramirez, J., & Wright, C. W. (1999). Sesquiterpene lactones from Neurolaena oaxacana. Journal of Natural Products, 62(8), 1093–1095. 10.1021/NP990038T/ASSET/IMAGES/LARGE/NP990038TF1.JPEG

Pramanik, P. K., Alam, M. N., Roy Chowdhury, D., & Chakraborti, T. (2019). Drug Resistance in Protozoan Parasites: An Incessant Wrestle for Survival. Journal of Global Antimicrobial Resistance, 18, 1–11. 10.1016/J.JGAR.2019.01.023

Radulović, N., Dekić, M., & Stojanović-Radić, Z. (2012). Chemical composition and antimicrobial activity of the volatile oils of Geranium sanguineum L. and G. robertianum L. (Geraniaceae). Medicinal Chemistry Research, 21(5), 601–615. 10.1007/S00044-011-9565-9/TABLES/3

Radulović, N. S., Stojković, M. B., Mitić, S. S., Randjelović, P. J., Ilić, I. R., Stojanović, N. M., & Stojanović-Radić, Z. Z. (2012). Exploitation of the antioxidant potential of Geranium macrorrhizum (geraniaceae): Hepatoprotective and antimicrobial activities. Natural Product Communications, 7(12), 1609–1614. 10.1177/1934578X1200701218;JOURNAL:JOURNAL:NPXA;CTYPE:STRING:JOURNAL

Riaz, A., Rasul, A., Kanwal, N., Hussain, G., Shah, M. A., Sarfraz, I., Ishfaq, R., Batool, R., Rukhsar, F., & Adem, Ş. (2020). Germacrone: A Potent Secondary Metabolite with Therapeutic Potential in Metabolic Diseases, Cancer and Viral Infections. Current Drug Metabolism, 21(14), 1079–1090. 10.2174/1389200221999200728144801/CITE/REFWORKS

Rodrigues, A. C. J., Carloto, A. C. M., Gonçalves, M. D., Concato, V. M., Detoni, M. B., Santos, Y. M. dos, Cruz, E. M. S., Madureira, M. B., Nunes, A. P., Pires, M. F. M. K., Santos, N. C., Marques, R.E. dos S., Bidoia, D. L., Borges Figueiredo, F., & Pavanelli, W. R. (2023). Exploring the leishmanicidal potential of terpenoids: a comprehensive review on mechanisms of cell death. Frontiers in Cellular and Infection Microbiology, 13, 1260448. 10.3389/FCIMB.2023.1260448/BIBTEX

Rodríguez-Chávez, J. L., Rufino-González, Y., Ponce-Macotela, M., & Delgado, G. (2015). In vitro activity of ‘Mexican Arnica’ Heterotheca inuloides Cass natural products and some derivatives against Giardia intestinalis. Parasitology, 142(4), 576–584. 10.1017/S0031182014001619

Sasidharan, S., & Saudagar, P. (2021). Leishmaniasis: where are we and where are we heading? Parasitology Research, 120(5), 1541–1554. 10.1007/S00436-021-07139-2/TABLES/3

Tang, Z. H., Liu, K. Y., Mei, J., & Gao, X. Z. (2010). [In vitro anti-Trichomonas vaginalis effects of a mixture of dihydroartemisinin and metronidazole]. Zhongguo Ji Sheng Chong Xue Yu Ji Sheng Chong Bing Za Zhi = Chinese Journal of Parasitology & Parasitic Diseases, 28(6), 416–421. https://europepmc.org/article/med/21500527

Tzanova, M., Grozeva, N., … M. G.-B., & 2024, undefined. (2024). Composition and antioxidant potential of essential oil of Geranium macrorrhizum L. from different regions of Bulgaria. Bcc.Bas.BgMT Tzanova, NH Grozeva, MA Gerdzhikova, MH TodorovaBULGARIAN CHEMICAL COMMUNICATIONS, 2024•bcc.Bas.Bg, 56, 2024. 10.34049/bcc.56.D.S1L11

World Health Organization (WHO). (2024). Leishmaniasis. https://www.who.int/data/gho/data/themes/topics/gho-ntd-leishmaniasis

Yu, H. W., Wright, C. W., Cai, Y., Yang, S. L., Phillipson, J. D., Kirby, G. C., & Warhurst, D. C. (1994). Antiprotozoal activities of Centipeda minima. Phytotherapy Research, 8(7), 436–438. 10.1002/PTR.2650080713

